## Supplementary Figures for "A temporal sequence of thalamic activity unfolds at transitions in behavioral arousal state"

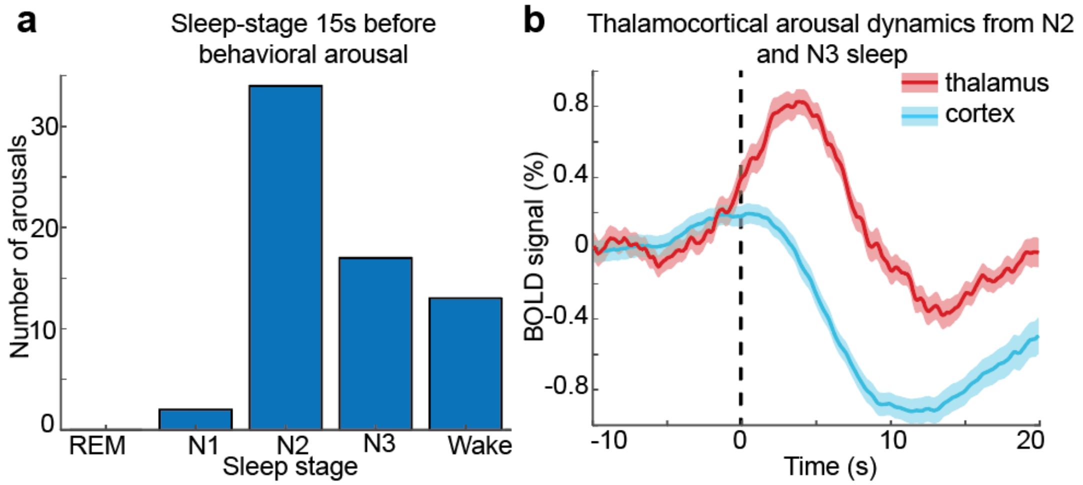


Supplementary Figure 1: a) Sleep stages 15 seconds before each arousal in Experiment 1 demonstrate that most arousals occurred from N2 and N3 sleep. b) When arousals from wake and N1 sleep are excluded, the same thalamocortical dynamics are preserved. Shading represents standard error.

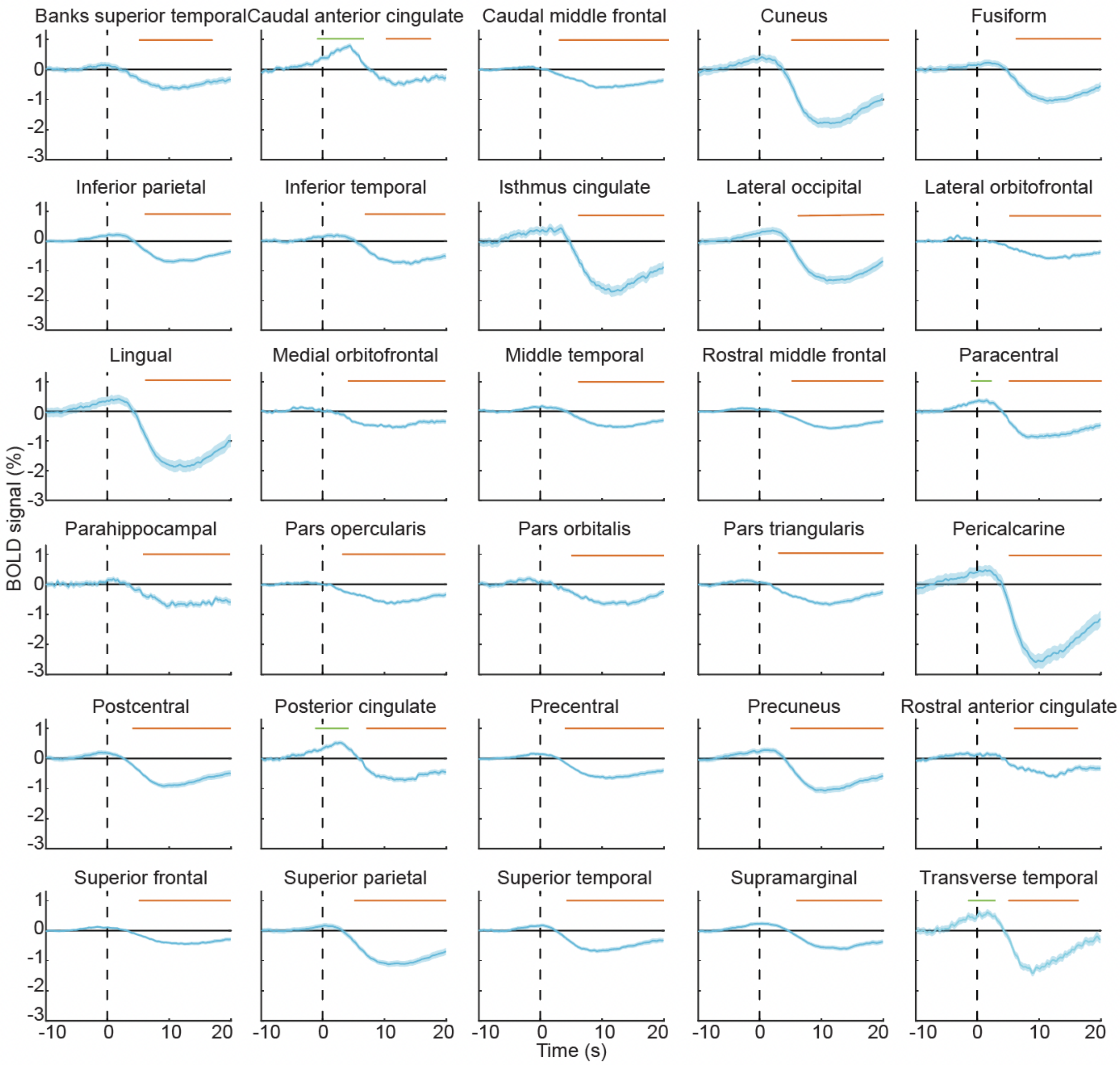


Supplementary Figure 2: All cortical regions at 3T during behavioral arousal. The vertical black dashed line represents the moment of behavioral arousal. A subset of cortical regions including the caudal anterior cingulate, posterior cingulate, paracentral, and transverse temporal cortices significantly increased during behavioral arousal (green bar). All cortical regions significantly decreased during arousal (orange bar). Shading represents standard error.

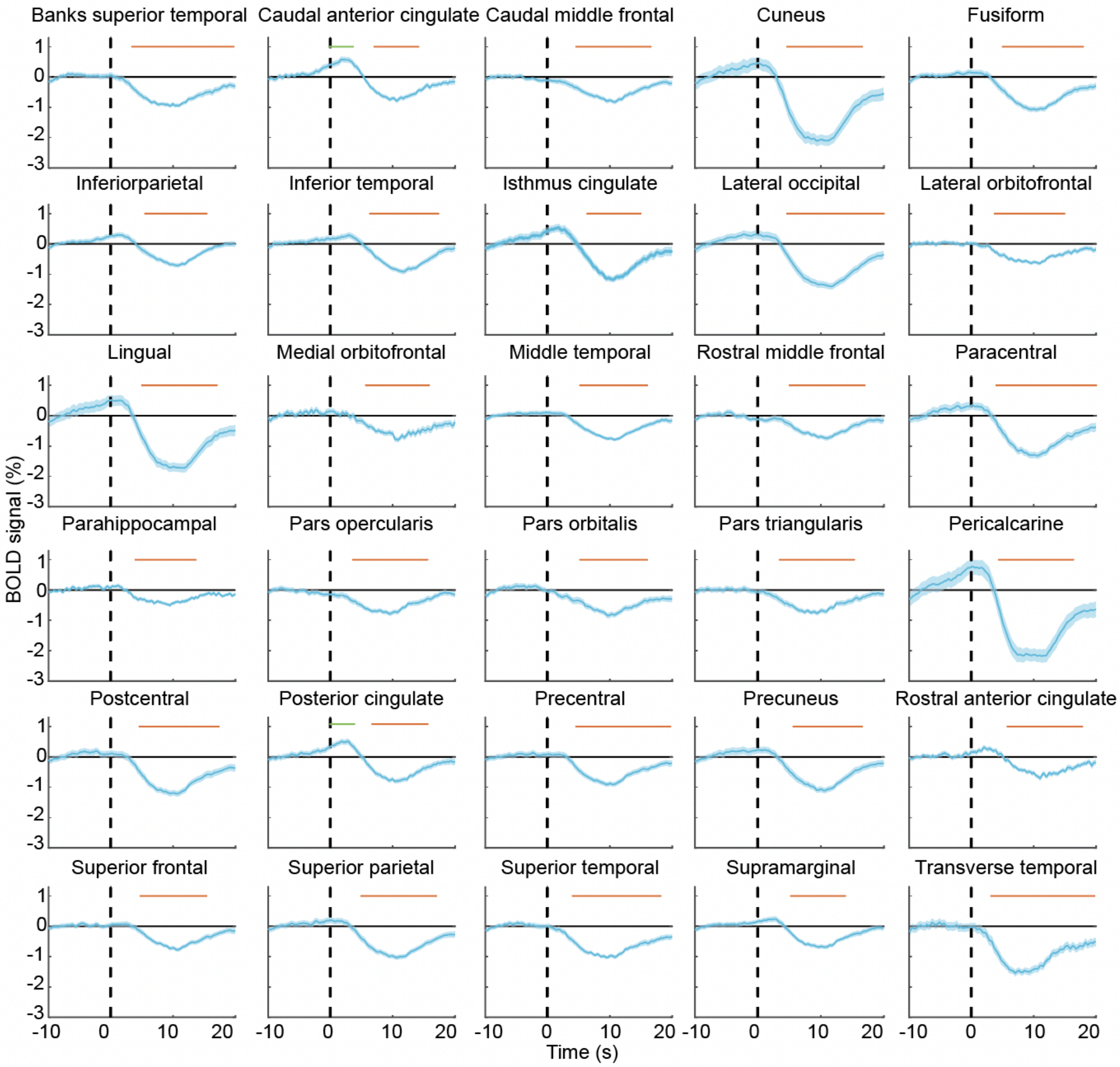


Supplementary Figure 3: All cortical regions at 7T during behavioral arousal. The vertical black dashed line represents the moment of behavioral arousal. A subset of cortical regions significantly increased during arousal (green bar), including the caudal anterior cingulate, and posterior cingulate. All cortical regions significantly decreased during arousal (orange bar). Shading represents standard error.

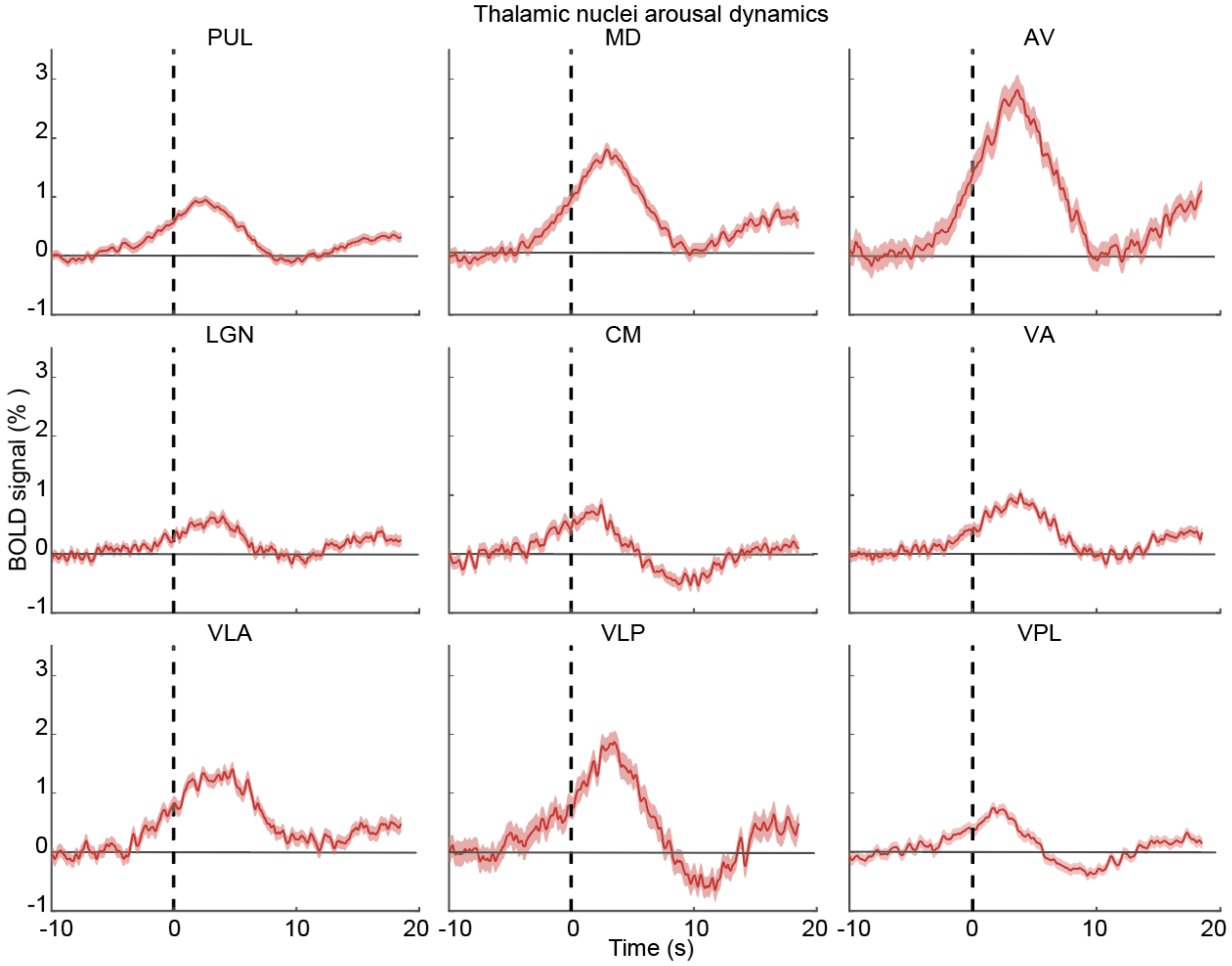


Supplementary Figure 4. Thalamic nuclei activate during behavioral arousal (vertical dashed line). Shading represents standard error.

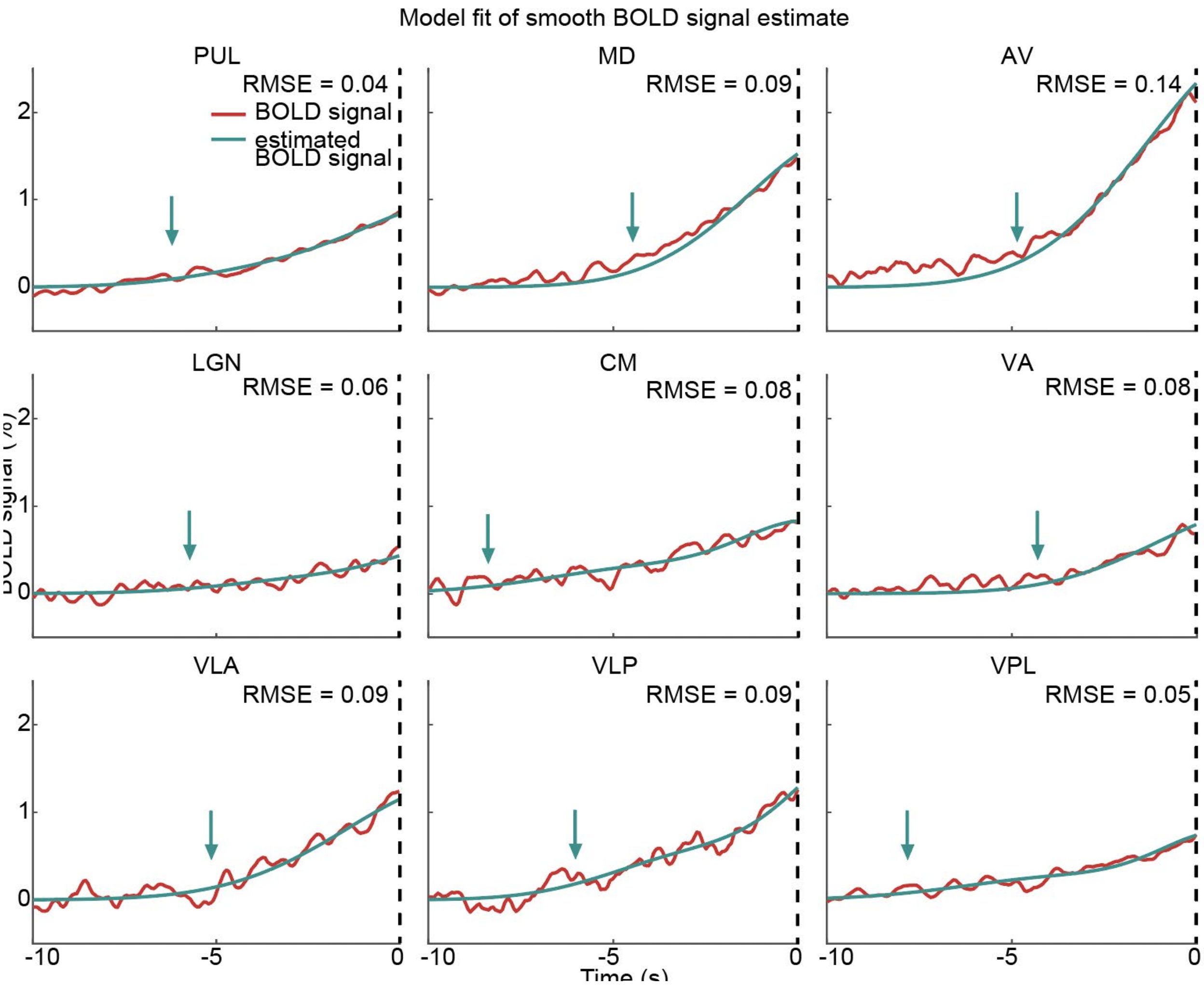


Supplementary Figure 5: Onset times of thalamic nuclei during behavioral arousal. The BOLD signal is in red and the model fit is in teal. The onset time of each thalamic nucleus is represented by the teal arrow. Behavioral arousal is marked by the black dashed line.

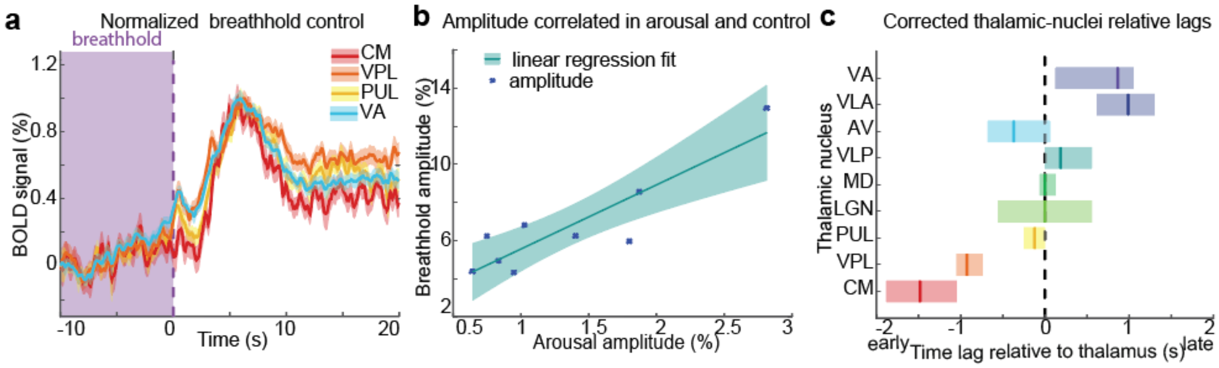


Supplementary Figure 6. Endogenous hemodynamics could not explain the observed thalamic nuclei sequence during behavioral arousal. a) Normalized signals from a subset of thalamic nuclei at breathhold release (purple dashed line). Shading represents standard error. b) The amplitude of the breathhold response across thalamic nuclei was highly correlated with their amplitudes during behavioral arousal, suggesting successful recapitulation of local SNR and hemodynamic properties. c) Correcting the thalamic sequence by subtracting the average lag during breathhold release does not greatly alter the activation sequence, with VPL and CM still showing earlier activity. The solid vertical bar represents lag time, and the shaded box is the 95% confidence interval. Color of each nucleus' lag represents order in original arousal sequence.

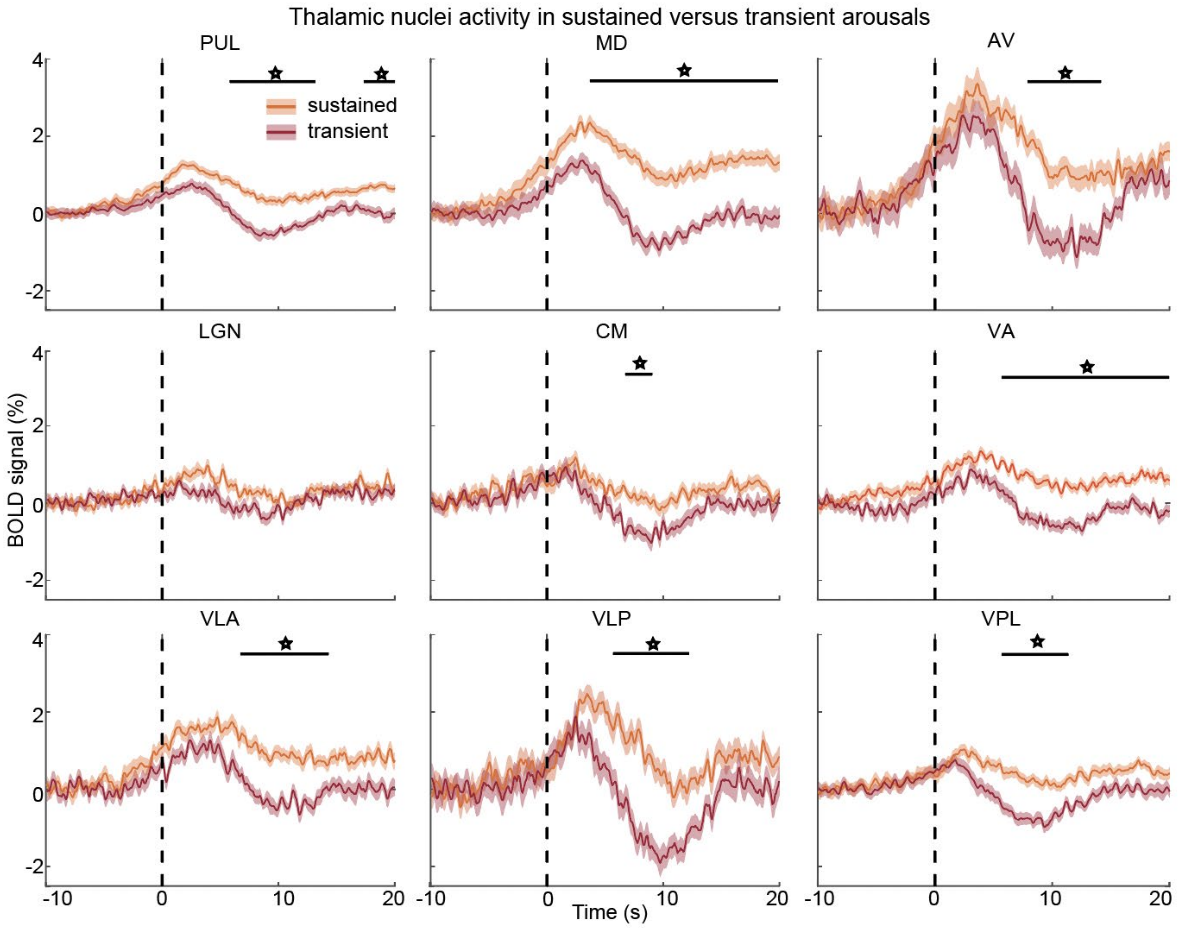


Supplementary figure 7: Thalamic activity at sustained vs transient arousals. Behavioral arousal is represented as the dashed vertical line. The response at sustained arousals is in orange, and at transient arousals is in red. Shading represents standard error. Most thalamic nuclei had significantly different post-arousal signals (starred horizontal line).

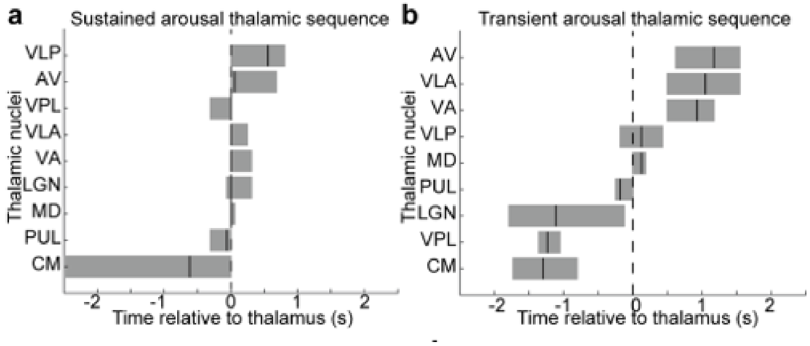


Supplementary figure 8: The activity sequence across thalamic nuclei differed in sustained vs transient arousals. a) The sequence at sustained arousals demonstrated that the activity across nuclei occurs closely together in time. Vertical line is the lag. Shaded rectangles represent standard error. b) The sequence at transient arousals demonstrated a larger lag between the early and late nuclei.
